# GOTHiC, a simple probabilistic model to resolve complex biases and to identify real interactions in Hi-C data

**DOI:** 10.1101/023317

**Authors:** Borbala Mifsud, Inigo Martincorena, Elodie Darbo, Robert Sugar, Stefan Schoenfelder, Peter Fraser, Nicholas M. Luscombe

## Abstract

Hi-C is one of the main methods for investigating spatial co-localisation of DNA in the nucleus. However, the raw sequencing data obtained from Hi-C experiments suffer from large biases and spurious contacts, making it difficult to identify true interactions. Existing methods use complex models to account for biases and do not provide a significance threshold for detecting interactions. Here we introduce a simple binomial probabilistic model that resolves complex biases and distinguishes between true and false interactions. The model corrects biases of known and unknown origin and yields a p-value for each interaction, providing a reliable threshold based on significance. We demonstrate this experimentally by testing the method against a random ligation dataset. Our method outperforms previous methods and provides a statistical framework for further data analysis, such as comparisons of Hi-C interactions between different conditions. GOTHiC is available as a BioConductor package (http://www.bioconductor.org/packages/release/bioc/html/GOTHiC.html).

## 2. Introduction

Hi-C is a high-throughput technique based on chromosome conformation capture to detect the spatial proximity between pairs of genomic loci^1,2^. It is now routinely used to study the three-dimensional folding of genomes^3-7^. In theory, a sequenced Hi-C read-pair should directly represent an interaction between two loci, with the number of mapped read-pairs corresponding to the frequency of interactions in the sample cell population. However, two challenges must be resolved in order to extract the true signal from Hi-C data.

The first is to identify and resolve systematic biases. Hi-C datasets present many effects common to high-throughput sequencing experiments, for instance amplification biases due to differences in sequence composition across the genome. There are also biases that are specific to Hi-C. For example, variations in the density of restriction sites cause large differences in genomic fragment sizes; since very long or very short fragments are difficult to ligate, both tend to be under-represented in the sequencing library^8^. The complex combination of known and unknown biases cause over- and underrepresentation of chromosomal regions when a Hi-C dataset is mapped to the reference genome. Thus, the number of observed read-pairs do not directly reflect the frequency of interactions between two genomic loci.

The second challenge is to distinguish between real and artefactual interactions. As depicted in Box 1A, a Hi-C library contains three types of read-pairs. (i) The first represents real interactions in which the ligation reaction occurs between the ends of a pair of crosslinked DNA fragments. (ii) The second corresponds to spurious self-ligations in which the ends of the same DNA fragment are ligated together; as the two ends of read-pairs map to the same DNA fragment, they are easily filtered. (iii) The third represents spurious ligations between two non-crosslinked DNA fragments; read-pairs from these reactions are problematic as they are indistinguishable from those arising through real interactions. The proportions of read-pairs representing real and spurious interactions can vary widely depending on the quality of the sample and library preparations.

**Box 1.**
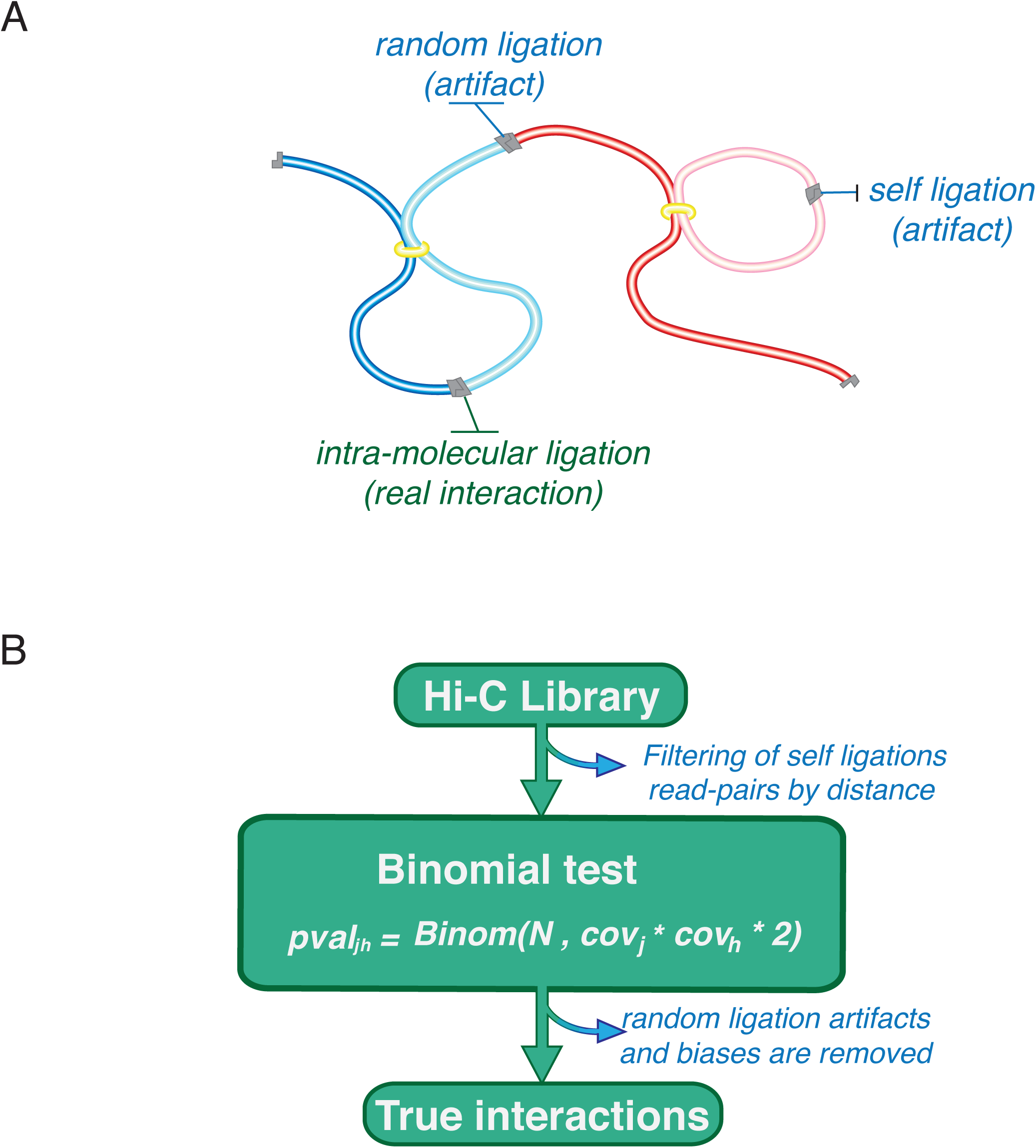
Schematic overview of the binomial model. (**A**) After crosslinking and digesting the chromatin, the DNA is ligated resulting in three types of ligation products. In order to detect real interactions, we first filter out self-ligations. With the remaining paired-reads, we then calculate the relative coverage across the genome in order to estimate the random interaction probability. (**B**) We finally apply the binomial test to distinguish between random and real interactions.

There are two main approaches to deal with biases. One approach is to identify the known sources of biases affecting observed read counts a priori, and model these biases. One such algorithm is *hicpipe,* which applies a multiplicative model to estimate the probabilities of interactions between two genomic regions as a function of mappability, fragment length and GC content; the numbers of mapped read-pairs are then normalised according to these estimates^8^. Another method, *HiCNorm,* models biases at lower resolution and uses Poisson regression for normalisation^9^. The other approach, used by most recent methods as well as us, is to assume that all biases are reflected in the observed read counts. An early example is *hiclib,* which proposes that the total bias is represented in the sequence coverage as the product of individual biases for each pair of genomic regions. Starting with the assumption that every genomic region should have identical coverages, *hiclib* iteratively normalises the original coverage until it becomes uniform along the whole genome^10^. This method has been implemented in a faster algorithm, *Hi-Corrector*^11^. *ChromoR,* also only uses the information encaptured in observed read counts. For normalisation of Hi-C data it uses Haar-Fisz Transformation to decompose the Poisson distributed read counts into Gaussian coefficients that are subsequently de-noised by wavelet shrinkage methods^12^.

Although both approaches have been frequently used, these methods exhibit several practical limitations. First, the assumptions behind the methods are untested against experimental control data and so their success in eliminating biases is unclear. Second, the issue of distinguishing between real and random interactions remains unresolved. Finally, the software implementation of some of these methods have either several dependencies that make them technically demanding to install and operate ^8,13^, require extensive pre-processing ^11,13^, or can not be applied at higher resolution ^12^.

Here, we introduce a straightforward binomial model that corrects the complex combination of known and unknown biases in Hi-C data (Box 1B). The model calculates accurately the probabilities that the observed number of read-pairs are due to random ligations, and yields a list of statistically significant interactions between pairs of genomic loci. We challenged the model using random ligation controls, demonstrating that the method gives high levels of specificity. GOTHiC, the accompanying BioConductor package, is fast, accurate and easy to use.

## 3. A binomial model for Hi-C data

For a given pair of genomic loci, GOTHiC calculates: (i) the probability of observing a given number of read-pairs between two loci through random ligations; and (ii) the effect size, “strength” or “frequency”, of interaction measured as the ratio of observed-over-expected numbers of interactions. GOTHiC assumes that the observed sequence coverage varies as a function of multiple known and unknown biases, including the density of restriction sites, cleavage efficiency, ligation efficiency, amplification and sequencing biases, and mappability. It assumes that the biases affect each end of read-pairs independently; thus the probability of observing a randomly occuring read-pair between two loci is modelled as the product of the relative coverages of the interacting loci. This is a reasonable assumption given our understanding of known biases^8-10^; the advantage of modelling the combined effect of biases is that it incorporates unknown sources and that it is robust against future variants of Hi-C methods.

First, self-ligations, dangling ends, re-ligations and incomplete digestion products are removed by filtering read-pairs mapping to the same fragment and within a specified distance of each other on the genome (default=10kb). Given the relative coverage of two genomic loci, *j* and *h*, the probability of a spurious read-pair linking the two loci can be calculated as:

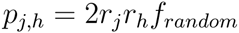

*r_j_*, the relative coverage of a locus, is calculated as:

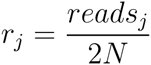

where *reads_j_* is the mapped read count for genomic locus *j,* and *N* is the total number of read-pairs in the filtered dataset. *f_random_* is the fraction of read-pairs in the Hi-C library arising from spurious ligations. Although *f_random_* could be estimated experimentally or computationally, in practise this may often be difficult and a conservative upperbound for *p_j,h_* can be obtained by excluding this term.

Given the probability of a read linking the two loci, the probability of observing *n* or more read-pairs between them by chance in a dataset of *N* reads, is given by the binomial cumulative density:

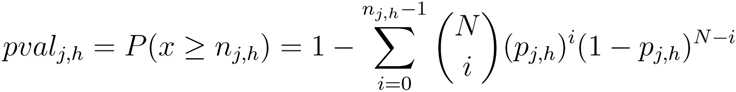

This yields a p-value for each interaction as a function of the coverage of both loci and the total number of reads in the experiment. Using the Benjamini-Hochberg multiple-testing correction (with L*(L-1)/2 tests, where L is the number of loci investigated), we obtain a q-value that can be used directly to identify statistically significant interactions at a pre-defined false discovery rate.

The log of the observed-over-expected ratio (R) can be used as a measure of effect size or as a normalised measure of interaction *frequency.*

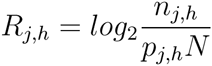

## 4. How well does GOTHiC perform?

### 4.1 Assessing performance using a Hi-C dataset and random ligation control

To assess performance, we applied GOTHiC at 1Mb resolution to two datasets generated from the same mouse fetal liver cell sample: (i) one produced using the standard Hi-C protocol and (ii) another containing only randomly ligated read-pairs with HindIII. The latter was produced by reversing the cross-links before the ligation step and it is analogous to an “input” control that is commonly used for background correction in ChlP-seq studies. As expected in a random control, 93–95% of read-pairs occur between loci on different chromosomes, in contrast to 20–40% of read-pairs in Hi-C datasets.

Read coverage is highly variable across the genome (Figure 1A): it correlates well with previously reported effects of GC content, mappability and restriction-site density, though not all variation is captured by these factors. The raw contact maps in Figure 1B emphasise how variations in sequence coverage affect the interpretation of unnormalised Hi-C data, in which regions of higher coverage ostensibly show stronger interactions and vice versa. Strikingly, the trend is apparent even in the random ligation control (blue arrow, right panel), which does not contain any true interactions. The high correlation in coverages between the real and random datasets (Pearson’s r=0.99) indicates that virtually all of the variation in coverage observed in a Hi-C sample is explained by experimental biases.

**Figure 1.**
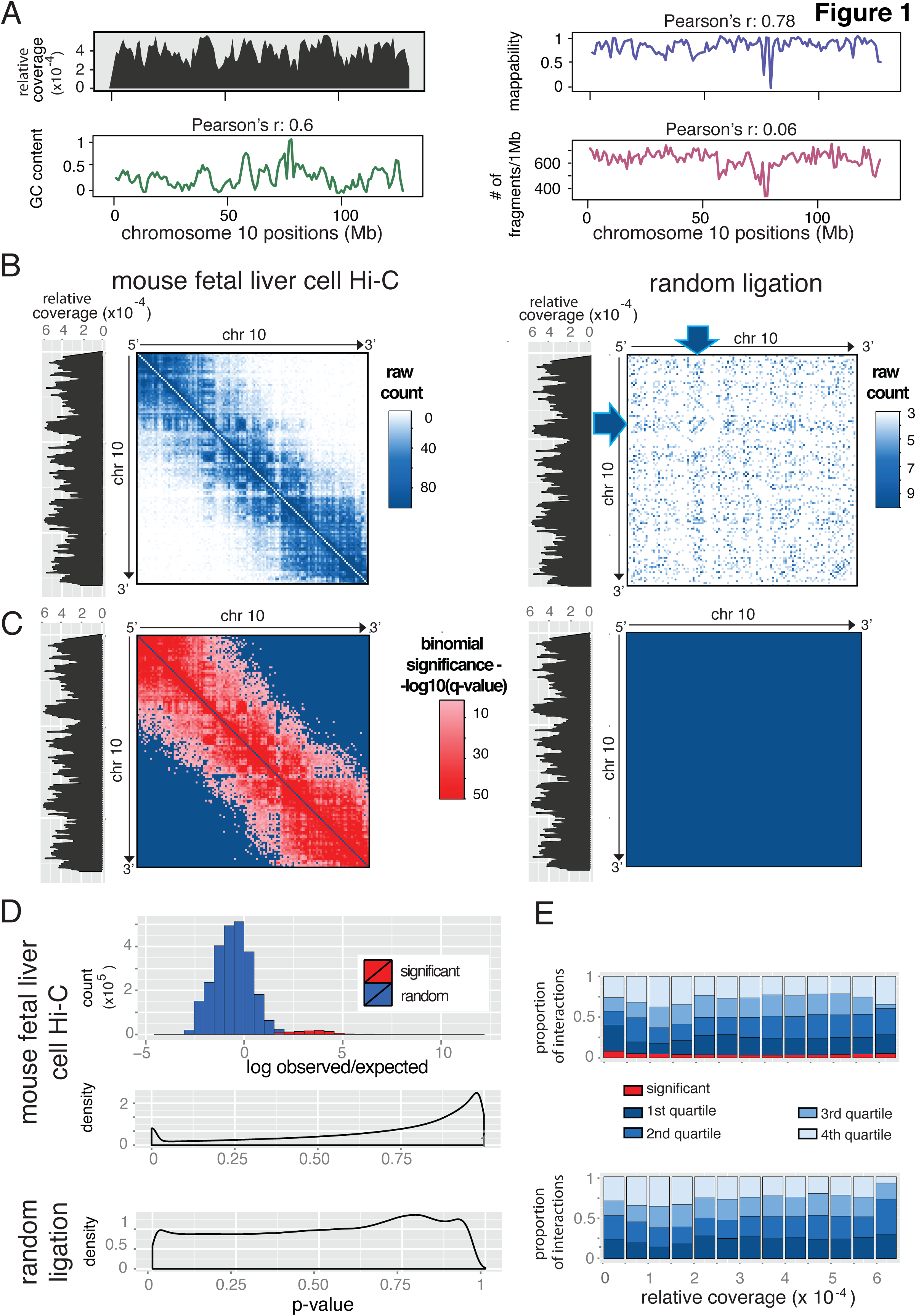
GOTHiC applied to mouse fetal liver Hi-C experiments. **(A)** From the top, distributions of the relative coverage, the GC content percentage, the mappability score and the number of fragments per 1Mb (y-axis) across mouse Chromosome 10 (x-axis in Mb) (GC content and mappability scores are as in^8^). **(B-C)** Contact maps of mouse Chromosome 10 containing raw read counts (interactions with at least 3 reads) and binomial significances respectively resulting from classic Hi-C experiment (left panel) and random ligation experiment (right panel) in fetal liver cells. The intensity of the signal is summarized by the gradient above each contact map. Significant interactions are colored with a red gradient in **C.** Arrows pinpoint a region of high coverage and its impact on the observed number of interactions (**B**, right panel). The coverage is represented at the left side of each contact map. **(D)** The top panel represents the distribution of observed/expected log ratio of significant (red) and non-significant (blue) interactions in the fetal liver cell sample. Middle and bottom panels represent the distribution of binomial p-values in the fetal liver cell and random samples respectively. **(E)** Influence of the relative coverage on the distribution of interaction significance. GOTHiC interaction ranking in the Hi-C (upper panel) and random ligation (lower panel) samples. The ranked lists were divided into quartiles, the first quartiles correspond to the top ranked interactions. Significant interactions are shown in red.

The processed contact maps in Figure 1C show how effectively GOTHiC deals with these biases, as the patterns influenced by underlying variations in coverage are removed (left panel). GOTHiC also identifies statistically significant interactions with high specificity (red squares, left panel). There is good separation in log (observed/expected) values between real and random interactions (Figure 1D, top), which is also reflected in the distribution of p-values (middle panel). GOTHiC identified ~90,000 statistically significant interactions in the Hi-C dataset (FDR <5%). In contrast, GOTHiC calls almost no interactions in the random ligation experiment (Figure 1C, right panel). This dataset confirms the specificity of the binomial model and the accuracy of FDR estimates, as violations of the underlying assumptions should lead to a large number of false positives. In fact, GOTHiC calls just 22 false positive interactions in the random ligation dataset from more than 3 million tests; this means that the p-values accurately reflect the probability of observing a given number of reads between any two loci as a result of experimental biases. This was also true at higher resolutions (500kb and 100kb bins) for these medium sized Hi-C experiments (~35M reads), as shown in Figure S1 and Supplementary Table 1.

In addition to calling statistically significant interactions, GOTHiC removes much of the underlying bias. Figure 1E demonstrates that the detection of significant interactions as well as the general ranking of interactions by their q-value is largely independent of coverage, as the proportion of significant interactions is stable across different coverage bins, and in each coverage bin the proportion of interactions falling into the different quartiles is near a quarter.

Alternatively to the q-value, the log-ratio, *R,* between the observed number of reads and the expected number of reads (log observed/expected) may be used as a normalised measure of interaction frequency. This value is similar to the log fold-change measure in differential expression analyses, and it would tend to show a high variance in regions of low coverage due to the low expected values, and the integer read counts, similarly to log fold-change of lowly expressed genes. However, the *R* value can be used for a dual cut-off to identify significant interactions above a desired effect size (as in volcano plots).

The output from GOTHiC can also be used to flag poor quality Hi-C libraries. We have observed that inadequate dilution or cross-linking can yield libraries with a high fraction of spurious read-pairs *(i.e.,* ligations between non-crosslinked fragments). As shown in the control dataset (Figure 1D), this will lead to a more uniform distribution of p-values, as expected by chance, and GOTHiC will successfully control the false discovery rate, yielding a small number of significant interactions.

### 4.2 Reproducibility between replicates using different restriction enzymes

It has been shown that treating the same biological sample with different restriction enzymes can cause large differences in coverage along the genome^3^. To evaluate the performance of GOTHiC in these conditions, we applied it at 1Mb resolution to previously published Hi-C datasets produced using HindIII and NcoI on a human lymphoblastoid cell line. These enzymes target distinct restriction motifs that are distributed differently along the genome; this results in different fragment densities, GC contents and mappability biases. Figure 2A highlights their remarkable impact on the coverage profiles and the raw contact maps (left and right panels, yellow highlighted boxes).

**Figure 2.**
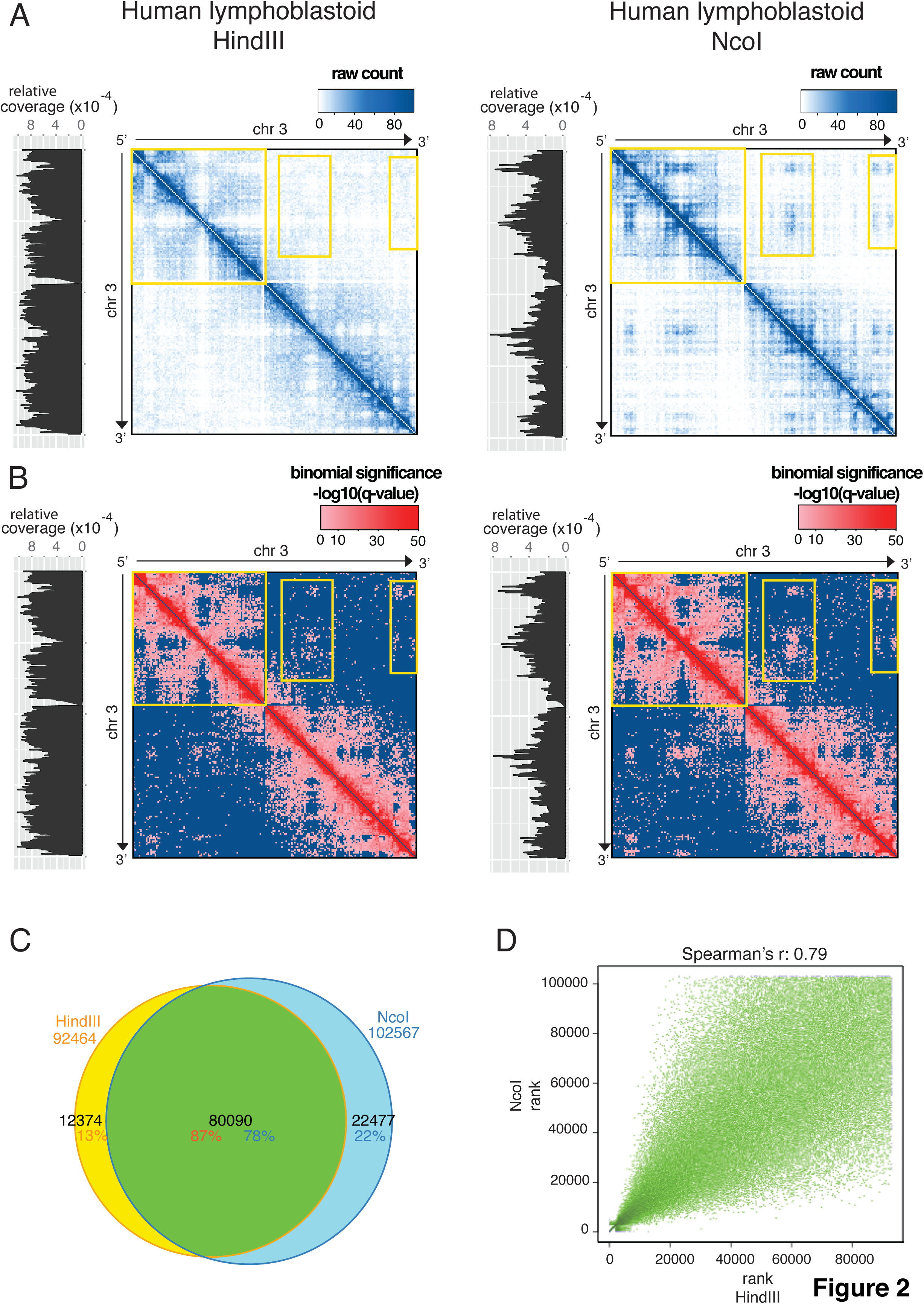
GOTHiC applied to human lymphoblastoid Hi-C experiments. **(A-B)** Contact maps of human Chromosome 3 containing raw read counts (interactions with at least 3 reads) and binomial significances respectively resulting from HindIII Hi-C experiment (left panel) and NcoI Hi-C experiment (right panel). The intensity of the signal is summarized by the gradient above each contact map. Significant interactions are colored with a red gradient in **B**. The coverage is represented at the left side of each contact map. **(C)** Venn diagram representing the overlap between significant interactions detected in HindIII (orange percentage) and NcoI (blue percentage) samples. **(D)** Correlation between the HindIII (x-axis)/NcoI (y-axis) common significant interactions (80,448 interactions) according to their rank. Spearman’s correlations are indicated above the plot.

Despite these strong biases, GOTHiC produces very consistent contact maps and statistically significant interactions (Figure 2B). Loci with very different numbers of read-pairs in the raw data are identified as interacting at similar significance levels after processing (Figure 2A and 2B, highlighted regions). We find 92,464 and 102,567 significant interactions in HindIII and NcoI experiments respectively, of which 80,090 overlap (Figure 2C), and the interaction rankings obtained from the two experiments show high correlation (Spearman’s r=0.79) (Figure 2D). The high overlap was maintained at higher resolutions (500kb, 100kb bins) (Figure S2, Supplementary Table 1).

## 5. Comparison with existing methods

Finally, in order to benchmark GOTHiC’s performance, we applied *hicpipe* and *hiclib,* which represent different normalisation approaches, to the mouse fetal liver and human lymphoblastoid Hi-C datasets (Figure 3, S3).

**Figure 3.**
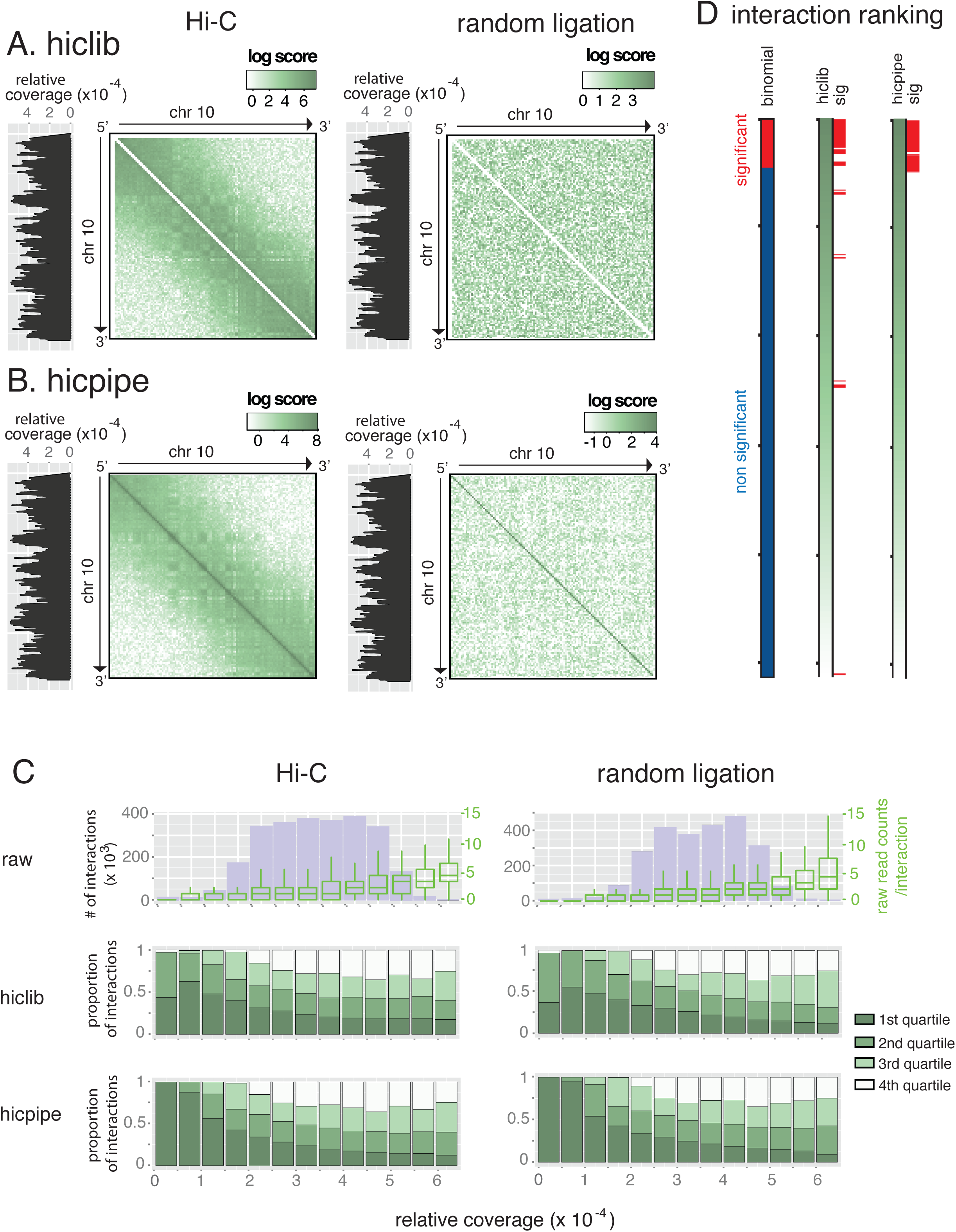
Comparison of mouse the fetal liver Hi-C data after processing by hiclib, hicpipe and GOTHiC. **(A-B)** Contact maps of mouse Chromosome 10 containing relative probability computed by hiclib and observed/expected log ratio obtained with hicpipe respectively resulting from classic Hi-C experiment (left panel) and random ligation experiment (right panel) in fetal liver. The intensity of the signal is summarized by the gradient above each contact map. **(C)** Influence of the relative coverage on the distribution of number of observed interactions (top panel), hiclib and hicpipe interaction ranking (middle and bottom panels), in the HiC (left) and random ligation (right) samples. The ranked lists were divided into quartiles, the first quartiles correspond to the top ranked interactions. The distribution of the number of reads per interaction is represented in the top panel with green box plots (corresponding y-axis is placed on the right of the plot). **(D)** Correspondence between binomial significant interactions (88292) and hiclib and hicpipe ranking. Blue bar corresponds to non-significant interactions from GOTHiC, red bar to significant ones. The green gradients represent the ranking of the interaction resulting from hiclib (left) and hicpipe (right) processing. Red bars indicate the significant interactions detected with GOTHiC.

As previously observed from the contact maps, the number of reads between two loci is strongly affected by the coverage of these loci (Figure 3C, boxplots in the top panel). Although the normalised interaction strength values from *hicpipe* and *hiclib* do not appear to show obvious biases in the contact maps (Figure 3A,B, S3A,B), more detailed assessment reveals that the outputs from both methods continue to suffer from coverage-dependent biases (Figure 3C, middle and bottom panels). The interaction strength measures are inversely correlated with coverage, suggesting overcorrection of the raw data - in other words, interactions in the 1^st^ and 2^nd^ quartiles for strength are enriched in the low coverage bins. In contrast, for GOTHiC both the significant interactions and the interactions ranked by q-values appear less affected by coverage (Figure 1E, Figure S3C b).

When comparing the overlap of significant or top-ranked interactions as detected by the three methods on both datasets, GOTHIC yielded the highest overlap and rank correlation, indicating a better removal of technical biases (Figures 2C,D, S3,D,E)

Finally, we examined the overlap of interaction scores between the three methods (Figure 3D, Figure S3F). Interactions identified as significant by GOTHiC tend to be highly ranked by *hiclib* and *hicpipe,* evidencing good agreement. Moreover, using the number of significant interactions returned by the binomial test of GOTHiC as a cut-off to select the top-ranked interactions returned by the other methods, revealed a very high overlap between all three methods (Figure S4A). However, we observed overcorrection of low coverage regions by *hicpipe* and hiclib. When comparing the coverage of interactions that were top-ranked by either of these methods but were not among the significant interactions by GOTHiC to those that were also called significant, we found that non-significant interactions had significantly lower coverage (two-tailed t test: p-value < 2.2e-16 for both *hicpipe* and *hiclib*) (Figure S4B). Thus, GOTHiC is at least as successful as existing methods in removing biases, but also provides significance values and a statistical framework for further analyses.

## 6. Discussion

Sequencing libraries produced by Hi-C experiments are noisy because of technical artifacts (self-ligations and random ligations) and complex biases caused by the intrinsic characteristics of the genome sequence (GC content, unequal distribution of restriction sites, uniqueness and mappability of the sequences). Here, we have proposed a simple solution to analyze Hi-C data using a simple binomial test, which successfully removes artifacts and sequencing biases to detect real genomic interactions even in the noisiest Hi-C datasets.

GOTHiC’s approach is simpler than existing methods, which require the identification and separate modeling of individual biases^8,9^, an iterative correction of biases^10,11^, or variance stabilisation ^12^. It yields similar rankings to previous methods, with comparable or even slightly improved bias removal and reproducibility between replicates. Most importantly, unlike any other method, GOTHiC calculates p-values that allow the identification of real genomic interactions and the removal of artefactual interactions with a well-controlled false discovery rate.

GOTHiC is implemented as an R package, which requires a mapped read file as input and returns a list of significant interactions. This implementation can analyze a whole-genome Hi-C dataset of 30 million uniquely mapped reads at 1Mb resolution in ~2 hours using a single core machine with ~200Mb memory, and can be several fold faster if run with the parallel option on more cores.

The sensitivity of the method could be further improved by estimating the fraction of inter-molecular ligations (*f_random_*). Our use of an upperbound (*f_random_*=1) provides a conservative estimate, ensures high specificity and should be preferred unless accurate information on the noise fraction across the genome is available.

Hi-C, as all 3C-type assays, captures both ‘regulatory/functional’ interactions (such as promoter-enhancer interactions) and ‘structural’ interactions, which are a consequence of overall higher-order folding of the chromatin fibre. Owing to structural reasons, neighbouring DNA regions are known to interact often, and HiC datasets typically show a near power-law decrease in interaction frequencies with increasing genomic distance. It is still a major challenge in chromatin biology to disentangle functional, structural, and functionally redundant interactions. One approach is to correct interaction frequencies by the expected frequency for a given genomic distance. This distance-correction assumes that functional interactions are stronger than other interactions at a given distance; however, the generality of this assumption is unclear. For example, it is likely that the strong structural interactions at close genomic distances ‘saturate’ the possible ligation-products in 3C-based assays, which would hinder the detection of regulatory interactions at short distances. Importantly, many enhancer-promoter interactions have been shown to act at relatively short distances^14^. To avoid relying on strong assumptions, GOTHiC does not perform a distance-correction, instead yielding a comprehensive list of biological interactions not explained by experimental noise. However, GOTHiC’s statistical framework can be modified such that the expected interaction frequencies are corrected for genomic distance, or alternatively q and R values can be adjusted after the identification of significant interactions as implemented by other methods^15^.

Finally, we envisage that the simple probabilistic framework introduced here could be further expanded to other applications in Hi-C, such as combining replicates, or identifying interaction changes between conditions. Significance levels and observed/exptected ratios obtained from GOTHiC can be used as the basis for algorithms predicting the 3D structure of genomes^16^ or those finding topologically associated domains^17^.

## 7. Materials and methods

### Tissue isolation

Fetal livers were dissected from C57BL/6 mouse embryos at day 14.5 (E14.5) of development. Fetal liver cells were filtered through a cell strainer (70 mm) and directly fixed in formaldehyde.

### Hi-C

Hi-C was performed essentially as described in Lieberman-Aiden et al.^3^, with some modifications. 30 to 50 million cells were fixed in 2 % formaldehyde for 10 min, quenched with 0.125 M glycine, spun down (400 x g, 5 min) and washed once with PBS. The cells were incubated in 50 ml permeabilisation buffer (10 mM Tris–HCl pH 8, 10 mM NaCl, 0.2 % Igepal CA-630, Complete EDTA-free protease inhibitor cocktail [Roche]) for 30 min on ice with occasional agitation, spun down (650 × g, 5 min, 4 °C), and the cell pellets were resuspended in 358 μl of 1.25 x NEBuffer2 (NEB) per 5 million cell aliquot. Eleven μl of 10 % SDS was added to each aliquot, followed by an incubation at 37 °C for 60 min with continuous agitation (950 rpm). To quench the SDS, 75 μl of 10 % Triton X-100 was then added per aliquot, followed by an incubation at 37 °C for 60 min with continuous agitation (950 rpm). To digest chromatin, 1500 U of Hind III (NEB) was added per aliquot and incubated at 37 °C overnight with continuous agitation (950 rpm). After digestion, restriction sites were filled in with Klenow (NEB) in the presence of biotin-14-dATP (Life Technologies), dCTP, dGTP and dTTP (all 30 μM) for 60 min at 37 °C. 86 μl of 10 % SDS was added per aliquot and incubated at 65 °C for 30 min with continuous agitation (950 rpm), followed by addition of 7.61 ml of ligation mix (745 μl of 10 % Triton X-100, 820 μl of 10 × T4 DNA ligase reaction buffer [NEB], 82 μl of 10 mg/ml BSA [NEB] and 5.965 ml water) per aliquot and incubation at 37 °C for 60 min with occasional agitation. For the ligation reaction 50 μl of 1 U/μl T4 DNA ligase (Life Technologies) was added per aliquot, followed by incubation at 16 °C for 4 h. The cross-links were reversed by adding 60 μl of 10 mg/ml proteinase K (Roche) per aliquot and incubating at 65 °C overnight. After overnight incubation, another 60 μl of proteinase K per aliquot was added, followed by incubation at 65 °C for an additional two hours. RNA was removed by adding 12.5 μl of 10 mg/ml RNase A (Roche) per aliquot and incubating at 37 °C for 60 min. DNA was isolated by a phenol (Sigma) extraction, followed by a phenol/chloroform/isoamylalcohol (Sigma) extraction and standard ethanol precipitation. The precipitated DNA was washed three times with 70 % ethanol, and dissolved in 25 μl TE per aliquot. Subsequently all aliquots were pooled and the Hi-C DNA was quantified (Quant-iT Pico Green, Life Technologies). Biotin was removed from non-ligated restriction fragment ends by incubating 30 to 40 μg of Hi-C library DNA with T4 DNA polymerase (NEB) for 4 h at 20 °C in the presence of dATP. After DNA purification (QIAquick PCR purification kit [Qiagen]) and sonication (Covaris E220), the sonicated DNA was end-repaired with T4 DNA polymerase, T4 DNA polynucleotide kinase, Klenow (all NEB) and dNTPs in 1 × T4 DNA ligase reaction buffer (NEB). Double size selection of DNA was performed using AMPure XP beads (Beckman Coulter), before dATP-addition with Klenow exo-(NEB). Biotin-marked ligation products were isolated with MyOne Streptavidin C1 Dynabeads (Life Technologies) in binding buffer (5 mM Tris pH8, 0.5 mM EDTA, 1 M NaCl) for 30 min at room temperature, followed by two washes in binding buffer, and one wash in 1 × T4 DNA ligase reaction buffer (NEB). PE adapters (Illumina) were ligated onto Hi-C ligation products bound to streptavidin beads for 2 h at room temperature (T4 DNA ligase in 1 × T4 DNA ligase reaction buffer [NEB], slowly rotating). After washes in wash buffer (5 mM Tris, 0.5 mM EDTA, 1 M NaCl, 0.05 % Tween-20) and binding buffer, the DNA-bound beads were resuspended in NEBuffer 2. Bead-bound Hi-C DNA was amplified with 12 PCR amplification cycles using PE PCR 1.0 and PE PCR 2.0 primers (Illumina). The concentration and size distribution of Hi-C library DNA after PCR amplification was determined by Bioanalyzer profiles (Agilent Technologies) and quantitative PCR, and the Hi-C libraries were paired-end sequenced on Illumina Genome Analyzer IIx.

## Publicly available data

Mouse random ligation sample: GSM1718028.

Human HindIII and NcoI lymphoblastoid Hi-C: GSE18199.

## Data access

Raw data have been submitted to the EBI ArrayExpress (https://www.ebi.ac.uk/arrayexpress/) under accession number E-MTAB-3891.

## 8. Figures

**Supplementary Figure S1.**
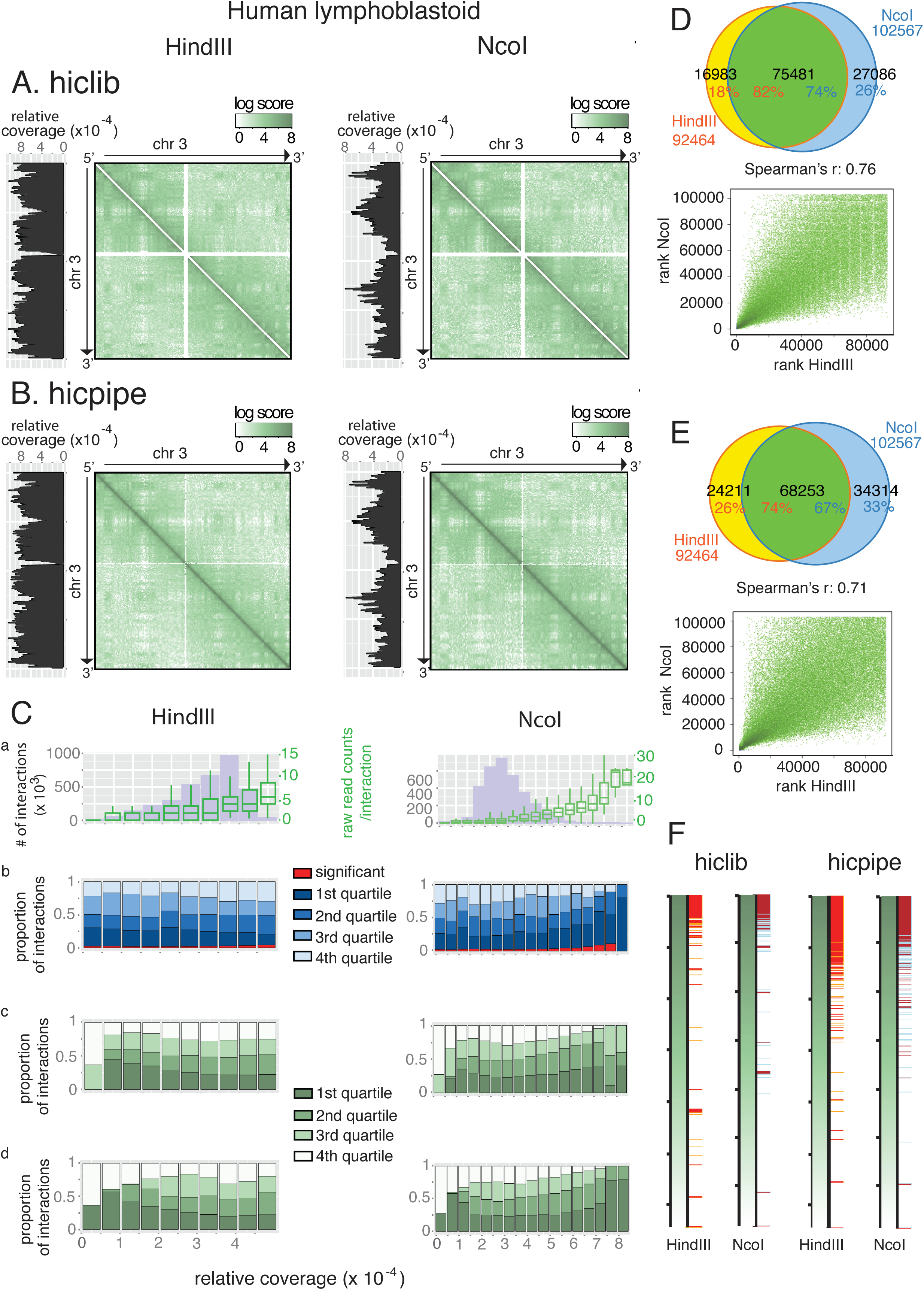
GOTHiC applied to mouse fetal liver Hi-C experiments at 500kb and 100kb resolutions. Contact maps of mouse Chromosome 10 representing binomial significances resulting from classic Hi-C experiment (upper panels) and random ligation experiment (lower panels) in fetal liver cells. Significant interactions are colored with a red gradient as on top.

**Supplementary Figure S2.**
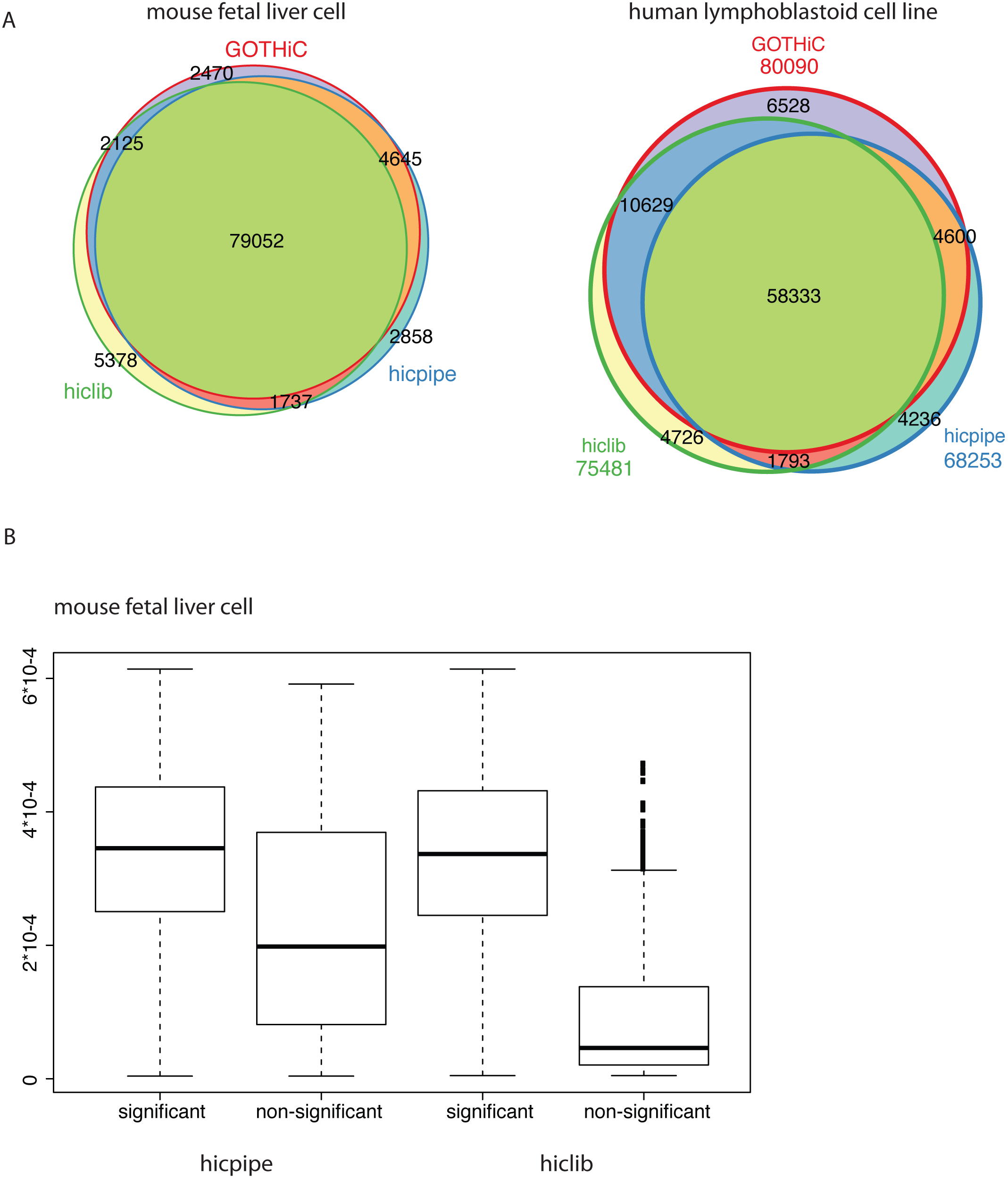
GOTHiC applied to human lymphoblastoid Hi-C experiments at 500kb and 100kb resolutions. Contact maps of human Chromosome 3 containing raw read counts representing binomial significances resulting from HindIII Hi-C experiment (left panels) and NcoI HiC experiment (right panels) at 500kb resolution (upper panels), 100kb resolution (middle and lower panels). The lower panels show a zoom in to chr3 1–10Mb. Significant interactions are colored with a red gradient as on top.

**Supplementary Figure S3.**
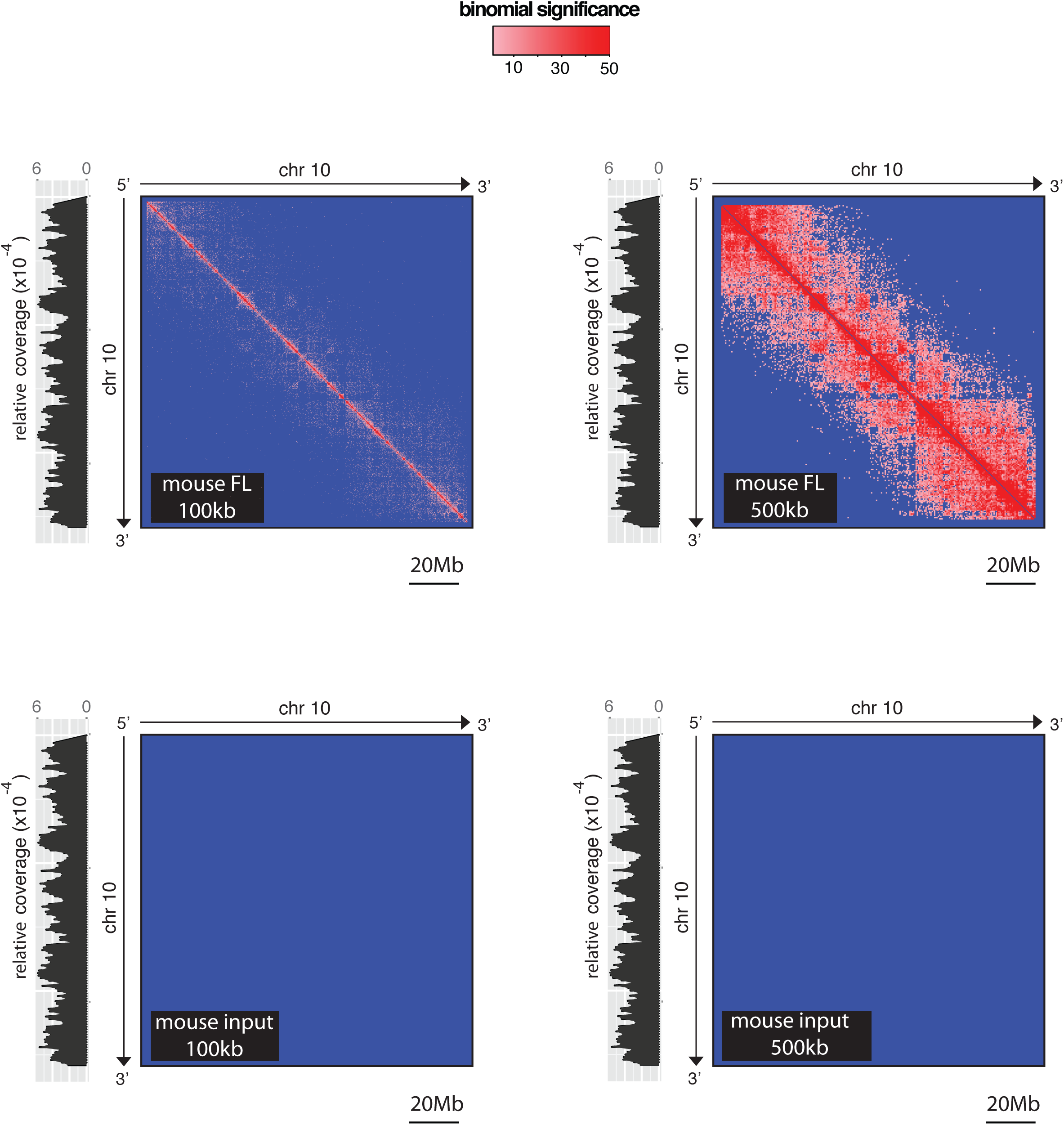
Comparison of human lymphoblastoid Hi-C data after processing by hiclib, hicpipe and the GOTHiC. **(A-B)** Contact maps of human Chromosome 3 containing relative probability computed by hiclib and observed/expected log ratio obtained with hicpipe respectively resulting from HindIII experiment (left panel) and NcoI experiment (right panel). The intensity of the signal is summarized by the gradient above each contact map. **(C)** Influence of the relative coverage on the distribution of (a) number of observed interactions, (b) GOTHiC, (c) hiclib and (d) hicpipe interaction ranking in the HindIII (left) and NcoI (right) samples. The ranked lists were divided into quartiles, the first quartiles correspond to the top ranked interactions. The distribution of the number of reads per interaction is represented in the top panel with green box plots (corresponding y-axis is placed on the right of the plot). 92,897 and 103,114 interactions were called significant using GOTHiC in the HindIII and NcoI samples respectively. In order to compare with the predictions of **(D)** hiclib and **(E)** hicpipe, we selected the 92,897 and 103,114 top ranked interactions of these methods and first computed the overlap (top) and correlation (bottom) between the two samples. **(F)** Correspondence between binomial significant interactions and hiclib and hicpipe ranking. The green-to-blue gradients represent the ranking of the interaction resulting from hiclib (left) and hicpipe (right) processing. Red bars indicate the significant interactions detected by GOTHiC in both HindIII and NcoI experiments. Orange bars indicate the significant interactions detected only in the HindIII experiment and blue bars indicate the significant interactions detected only in the NcoI experiment.

**Supplementary Figure S4.**
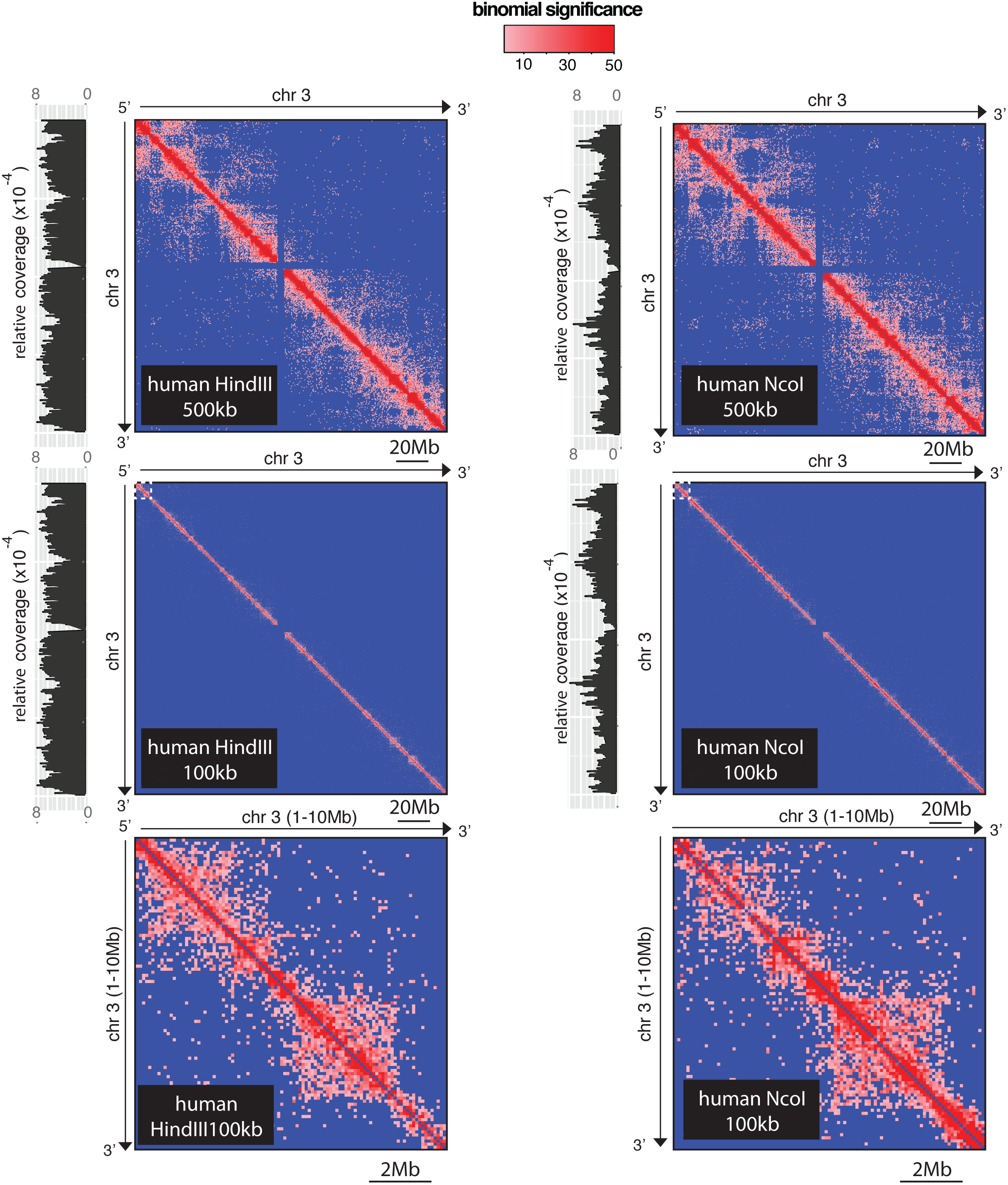
Overlap of top-ranked interactions from hiclib and hicpipe with significant interactions from GOTHiC. GOTHiC identified 88,292 significant interactions in the mouse fetal liver cell Hi-C dataset. **(A)** Venn diagram showing the overlap between the significant interactions identified by GOTHiC and the top 88,292 interactions from the hiclib and hicpipe outputs. **(B)** There were 80,448 significant interactions detected by GOTHiC that overlapped between the HindIII and NcoI experiments in the human lymphoblastoid cell line. The Venn diagram shows the overlap of between the GOTHiC, hiclib and hicpipe outputs.

**Supplementary Table 1.** Number of significant interactions in mouse fetal liver and human lymphoblastoid cells identified by GOTHiC at higher resolutions.

| # of significant interactions | 100kb | 500kb | 1Mb |
| --- | --- | --- | --- |
| input | 745 | 25 | 22 |
| fetal liver | 448070 | 246596 | 88299 |
| NcoI | 360516 | 192492 | 102567 |
| HindIII | 295378 | 159888 | 92464 |
| overlapping | 182038 | 117181 | 80090 |

